# Genomic epidemiology and phenotypic characterisation of *Salmonella enterica* serovar Panama in Victoria, Australia

**DOI:** 10.1101/2024.07.23.603842

**Authors:** Samriddhi Thakur, Sarah L. Baines, Cheryll M. Sia, Mary Valcanis, Louise Judd, Benjamin P. Howden, Hayley J. Newton, Danielle J. Ingle

## Abstract

*Salmonella enterica* serovar Panama, a causative agent of non-typhoidal salmonellosis (NTS), is one of several serovars that causes invasive NTS disease (iNTS) in humans. *S.* Panama is an understudied pathogen, with its pathobiology poorly understood. It is a predominant iNTS serovar in Australia, a high-income country with high rates of salmonellosis, where *S.* Panama has been documented to have a high odds ratio for causing iNTS. This study investigates the genomic epidemiology and antimicrobial resistance profiles of all *S.* Panama isolates recovered in Victoria, Australia, between 2000 and 2020. We examined the infection dynamics of *S.* Panama in seven isolates, representing the genetic diversity of the study population. Two sub-lineages, encompassed within a previously described Asian lineage, were identified. Multi-drug resistance (resistance to ≥3 drug classes) was detected in 46 (51.7%) Australian isolates. The plasmid-mediated colistin resistance gene, *mcr1.1*, was detected in one Australian *S.* Panama isolate, carried by an IncI plasmid previously reported in *Salmonella* and *Escherichia coli* isolates collected from poultry in South-East Asia. Examination of the intracellular replication dynamics of *S.* Panama isolates demonstrated diverse phenotypes. In THP-1 derived macrophages, despite low host cell uptake, *S.* Panama showed higher replication rates over time compared to *S. enterica* serovar Typhimurium. However, a causative genotype could not be identified to explain this observed phenotype. This study provides insights into the *S.* Panama isolates imported into Australia over two-decades, showing MDR was common in this iNTS serovar, and colistin resistance reported for the first time. It provides the first data on the host-pathogen interactions of *S.* Panama in Australia, which will aid our collective understanding of the pathobiology of *S.* Panama and iNTS serovars more broadly.

**Author Summary:** In Australia, non-typhoidal *Salmonella* (NTS) cases have been on the rise since the 1970s; characterised by self-limiting enteritis, some NTS infections can result in systemic infections called invasive NTS disease. *Salmonella enterica* serovar Panama is a leading iNTS serovar in Australia. This study characterised the genomic epidemiology of *S.* Panama, identifying two lineages circulating in Australia over two decades and placing them within a global context. It also investigated the antimicrobial resistance (AMR) mechanisms of *S.* Panama, with multi-drug resistance commonly observed. Further, it identified the first plasmid-mediated colistin-resistant *S.* Panama in Australia. We additionally examined the characteristics of *S.* Panama-mediated host-pathogen interactions in both epithelial and macrophage cells lines, providing the first insight into the infection dynamics of this understudied pathogen. Thus, this study combines genomics and *in vitro* infection experiments to understand the pathogenic behaviour of the neglected iNTS *S.* Panama.

## Introduction

Non-typhoidal *Salmonella* (NTS) consists of over 2,600 serovars and commonly causes self-limiting gastroenteritis (1–3). It is estimated that NTS infections cause a disease burden of 94 million cases with 155,000 deaths per annum (2). A subset of NTS serovars cause systemic disease with prolonged febrile symptoms. These are known as invasive non-typhoidal *Salmonella* (iNTS) (4), with 77,500 deaths attributed to iNTS globally each year (5). In human disease, the invasive potential can differ across NTS serovars or even between lineages of a serovar (6–8). Examples of iNTS serovars include *S.* Typhimurium, *S.* Enteritis, *S*. Choleraesuis, *S*. Dublin, *S*. Schwarzengrund, *S*. Heidelberg, *S*. Virchow, *S*. Infantis, and *S*. Panama (9). Previous studies have largely focussed on *S*. Typhimurium, *S*. Enteritidis and *S*. Dublin (10–16). For example, investigation into the genetic mechanisms associated with pathogenicity of *S*. Typhimurium ST313, which is the leading iNTS serovar in sub-Saharan Africa (17–20). However, the genomic epidemiology and host-pathogen interactions of other iNTS serovars, including *S.* Panama, have not been well characterised to date (21–23).

*Salmonella* Panama is an iNTS serovar with a broad host range, associated with bloodborne infections and meningitis in both children and adults (23). In high-income countries (HICs), *S.* Panama has frequently been epidemiologically linked to foodborne infections in humans via pork derived food products and international travel (24–26). Antimicrobial resistance (AMR) is an emerging issue in *S.* Panama, with multidrug resistance (MDR; resistance to ≥3 drug classes) detected in isolates linked to both Europe and Asia (22, 27, 28). The global population structure of *S.* Panama was recently described, defining multiple lineages with strong geographical associations; IncN plasmids were identified as a likely source for the spread of MDR in recent European isolates. *S*. Panama isolates collected in Australia were found to form a sub-lineage with isolates from Asia, suggestive of an established lineage within the broader geographical region (22).

In Australia, *S*. Panama has been identified in epidemiological studies as one of the leading iNTS serovars with a high odds ratio for causing invasive infections (35, 37). However, a detailed genomic investigation of *S.* Panama circulating over Australia has yet to be undertaken. Moreover, the host-pathogen interactions that mediate *S.* Panama infection are poorly understood, with no prior study examining the pathobiology of this serovar. Unbiased whole genome sequencing (WGS) of all cultured *S.* Panama isolates provides the opportunity to better understand the genomic epidemiology and evolution of this understudied pathogen. Further, WGS based surveillance methods can be utilised to monitor the emergence of AMR, a well-established global threat to public health (29, 30). Accordingly, we performed WGS on all cultured *S.* Panama isolates in the state of Victoria, Australia over the past two decades to better understand the *S.* Panama circulating over this period, and to characterise the AMR mechanisms and plasmids within the population. We combined genomic data with source of collection to infer the type of infection (invasive or gastrointestinal disease), to explore the *in vitro* infection dynamics of the serovar.

## Methods

### Data Source for Australian *S*. Panama isolates

The Microbiological Diagnostic Unit Public Health Laboratory (MDU PHL) is the bacterial reference laboratory for the state of Victoria, Australia. At MDU PHL *Salmonella* isolates were phenotypically characterised to determine serovars until the implementation of routine WGS in 2018. For this study, the WGS sequencing data was generated for a total of 92 *S. enterica* samples, collected from human faecal or invasive (non-faecal) sources between 2000 and 2021. Duplicate *S.* Panama isolates collected from the same patient within two weeks were considered to be the same infection and were excluded from this study. Data were collected in accordance with the Victorian Public Health and Wellbeing Act 2008. Ethical approval was received from the University of Melbourne Human Research Ethics Committee (study number 1954615.3).

### Isolate selection strategies for construction of global phylogeny

The rationale for including publicly available WGS data was to place the *S*. Panama isolates collected in Australia with *S.* Panama circulating globally. Paired end reads of *S*. Panama isolates were downloaded from the European Nucleotide Archive (ENA) browser (https://www.ebi.ac.uk/ena/browser/home). Only those from human infections recorded between 2000 and 2021 were considered, since the Australian isolates in the current study were collected in the same timeframe. The selection strategies for each public BioProject were as follows:

i. PRJNA248792 (UK): Most isolates included in this BioProject belonged to multi-locus sequence type (ST) 48, with some exceptions (single-locus variants of ST48 and novel multi-locus variants (MLV) of ST48). All these non-ST48 isolates were initially selected to include diversity in the dataset, with the novel MLV isolates removed after quality control. For the ST48 isolates, all isolates from a particular year were considered if the year recorded ≤5 isolates, otherwise five isolates/year were randomly sampled.
ii. PRJNA230403 (USA): Only those isolates that were sequenced by the Centers for Disease Control (CDC) were considered. All isolates from a year were considered if the total number was ≤5; if >5, random sampling of 5 isolates/year was performed.
iii. PRJNA543337 (Canada): Public Health Canada only recorded isolates from years 2015 and 2016 in this BioProject; 5 isolates/year were randomly selected from each year.
iv. PRJEB18998 (Ireland): All five *S*. Panama isolates were considered.
v. PRJNA203445 (USA): One *S*. Panama isolate from Thailand was considered for the global dataset.

### DNA extraction and short read whole-genome sequencing

DNA extraction was performed as described by Sia *et al.,* followed by generation of 2x150 bp paired-end reads using Illumina NextSeq500/550 (31).

### Quality controls and draft genome assembly

For the Illumina reads of each isolate, three QC parameters were considered: (i) average read depth of >50, (ii) ≥50% of reads per isolate with Phred Score of Q ≥ 30, and (iii) taxonomic assignment identified ≥40% of ƙ-mers as belonging to *Salmonella enterica* using Kraken v2.1.2 (32). Draft genome assemblies were generated for all Australian and public *S*. Panama isolates using SPAdes v3.15.2 (33) with default parameters; assemblies resulting in >500 contigs were excluded from the dataset.

### MLST and *in silico* serovar determination

*In silico* multi-locus sequence typing (MLST) was performed on all local and public draft genomes with mlst v2.23.0 (https://github.com/tseemann/mlst), using the pubmlst database ‘*senterica’*. The *Salmonella In Silico* Typing Resource tool, SISTR v1.1.1 was used to confirm the serovars from the draft assemblies of both Australian and public isolates (34).

### Inferring the global maximum likelihood phylogeny

For inferring the global phylogeny, 89 Australian *S*. Panama isolates (that satisfied QC parameters and were confirmed to be of the correct serovar *in silico*) were considered. Similarly, 73 public isolates were considered from the following countries: United Kingdom (n=40), United States of America (n=18), Canada (n=9), Ireland (n=5) and Thailand (n=1). *S.* Panama ATCC 7378 (CP012346.1) was used as the reference strain since this was the only publicly available complete genome for the serovar at the commencement of this study. *S.* Panama ATCC 7378 was isolated in the USA in 2013. The total 162 isolates were aligned to the reference genome using Snippy v4.4.5 (https://github.com/tseemann/snippy) with default parameters. Coordinates of phage regions were detected using the Phaster webtool (https://phaster.ca/), with identified phage regions masked during construction of a core genome alignment (35, 36). Regions of recombination were identified using Gubbins v3.2.1 and removed to generate the final alignment for phylogeny construction (37). IQ tree v2.1.4 was used to construct the serovar-level maximum likelihood (ML) tree, using a general time reversible (GTR) substitution model +Γ, inclusive of invariable constant sites and 1000 ultra-fast bootstrap replicates (38). Bayesian Analysis of Population Structure (BAPS), a probability-based algorithm for clustering genetically similar subpopulations, was run with 2 levels and 20 initial clusters to provide statistical confidence for lineage identification using the R package RHierBAPS (39, 40).

### Inferring the local *S*. Panama phylogeny and estimating pairwise SNP distances

80 Australian isolates belonging to lineages BAPS 3 and BAPS 4 were mapped to a complete local reference isolate AUSMDU00067783, generated in this study (see selection of isolates and long read sequencing methods below), using Snippy v4.4.5 (https://github.com/tseemann/snippy). The workflow for inferring the ML phylogeny was the same as described previously. The core-genome alignment file for the local phylogeny was further used to determine the pairwise SNP-distances using the tool snp-dists v0.8.2 (https://github.com/tseemann/snp-dists) and was visualized with the ggplot2 v3.4.4 (41) package in RStudio v22.12.0 (42).

The significance level for association of BAPS 3 and BAPS 4 with isolate source was determined using GraphPad Fischer’s exact test (two-tailed P value) online calculator (https://www.graphpad.com/quickcalcs/contingency1/).

### Characterisation of the AMR determinants

Genomes were screened for known AMR determinants using AbriTAMR v1.0.14 (29), with NCBI’s AMR Finder Plus database v3.10.16 (43), and including the species parameter “*Salmonella*” to detect species specific point mutations. The intrinsic efflux resistance genes *mdsA* and *mdsB* were detected *in silico* in all isolates (Table S1) and not considered for analyses. The AMR profiles for the draft assemblies were visualised with the R package ComplexUpset v1.3.3 (44, 45).

### Detection of plasmid replicons

Draft genome assemblies were screened for known plasmid replicons using the PlasmidFinder database (46, 47) in ABRicate v1.0.1 (https://github.com/tseemann/abricate), with a minimum percentage nucleotide identity of 90% and minimum coverage of 90%.

### Isolate selection strategies for long read sequencing and phenotypic characterisation

Considering AMR profiles, source (faecal or blood culture), population structure and year of isolation, seven isolates were selected for long read sequencing with Oxford Nanopore Technologies (ONT). Two pairs from BAPS 3 and one pair from BAPS 4 were selected with respective AMR profiles, along with a colistin resistant invasive isolate from BAPS 3. For selecting matched faecal and invasive pairs, AMR gene profiles were first matched between each pair, with preference given to same year of isolation and sharing of the same immediate common ancestor. The latter two criteria were relaxed for BAPS 4 isolate pair. Further details of the selected isolates are provided in (Table S1).

### Long read sequencing and complete genome assembly

The isolates were cultured overnight at 37°C Luria-Bertani (LB) Miller agar, and DNA was extracted using GenFind V3 according to the manufacturer’s instructions (Beckman Coulter) using lysozyme and proteinase K without size selection. Sequencing libraries were made with the SQK-NBD112.96 kit and sequenced on R10 MinION flow cells. Base-calling of the long reads was performed using Guppy v.3.2.4, the inbuilt super accuracy base-caller of Nanopore GridION X5. The approach for processing the ONT long read data was based upon Baines *et al.* 2020 (48). Porechop v0.2.4 (https://github.com/rrwick/Porechop) was used to perform adaptor-trimming and demultiplexing, using default parameters except a minimum identity percentage of 85% for barcode identification. The resulting long-reads were then filtered using Filtlong v0.2.1 (https://github.com/rrwick/Filtlong) with the following parameters: i) reads <1000 bp were excluded; ii) during read scoring, read quality and read length were given equal weighting, and; iii) a maximum of 460,000,000 bases were kept (equivalent to 100-fold coverage for a 4.6 Mb genome).

Genome assembly was performed using the Dragonflye pipeline v1.0.13 (https://github.com/rpetit3/dragonflye) with default parameters, and one round of short-read error correction with Polypolish v.0.5.0 (49). The genomes were annotated using Bakta v.1.6.1, specifying the genus as *Salmonella* and setting a minimum contig length of 200 bp (50).

### Comparison of MDR regions in complete genomes

The MDR regions of interest were extracted as Genbank files using Geneious Prime v2023.1.1 (https://www.geneious.com) from the annotated assemblies. Easyfig v2.2.5 was then used to compare the similar gene regions and output pairwise alignment files (51). These were then plotted in R using the gggenomes v0.9.5.900 package to show the similarities between genetic regions (52).

### Characterisation of plasmid with colistin resistance mechanisms

To identify similar publicly available plasmids with colistin resistance mechanisms, the plasmid sequence of AUSMDU00067711 was screened against the NCBI database (Nt) with blastn (https://blast.ncbi.nlm.nih.gov/Blast.cgi). The top four hits were selected (AP018355.1, OM038692.1, MN232197.1 and KU934209.1) which all had >99% query coverage and >90% identity. MN232197.1 was annotated for this study using Bakta v.1.6.1 (50). To visualise the comparison of the identity of *mcr-1.1* gene containing IncI2 plasmid from AUSMDU00067711 to public reference plasmids Blast Ring Image Generator (BRIG) v0.95 was used (53).

### SNP and pangenome comparison of closely related faecal-invasive isolate pairs

Snippy v4.4.5 (https://github.com/tseemann/snippy) was used to identify SNPs, insertions and deletions between closely related faecal and invasive pairs. For the BAPS 3 pair, paired end reads of AUSMDU00067873 were mapped to the complete genome of AUSMDU00068128. For the BAPS 4 pair, AUSMDU00068423 was used as reference to map AUSMDU00068404 reads. To compare the gene content between the pairs in each BAPS group, the complete genomes of all the four isolates were annotated with Prokka v1.14.6 (55). To investigate gene content between individual pairs, Panaroo v1.3.3 (56) was used with default parameters in conservative mode. Genes of interest were further characterised using *blastp* on NCBI, with NCBI’s Conserved Domains Database (57, 58).

### Bacterial isolates and growth conditions

The seven bacterial isolates selected for ONT sequencing were also phenotypically characterised for bacterial replication (Table S1). The *S*. Panama isolates and the *S*. Typhimurium SL1344 reference strain were streaked on Luria-Bertani (LB) Miller agar and stored at 4°C for up to two weeks. The *S*. Panama isolate AUSMDU00068404 was consistently observed to have two distinct colony morphologies. One colony type was observed to be larger and irregular (AUSMDU00068404 LCV) than the other (AUSMDU00068404). Overnight cultures were grown by inoculating a single colony in 3ml of LB Miller broth and incubating at 37°C under shaking condition of 180 rpm. For inducing type 3 secretion system 1 (T3SS1) expression, the protocol described by Klein *et al.* 2017 was followed with minor modifications (54). Briefly, 3% v/v overnight cultures were sub-cultured in 10ml LB Miller broth at 37°C, 200 rpm for 3.5 hours, then used as the inoculum for the *in vitro* infection experiments.

### Phenotypic comparison of bacterial replication in epithelial and macrophage cell lines

The *S*. Typhimurium SL1344 type strain was included as a non-invasive control strain for comparing bacterial internalisation and survival in both epithelial and macrophage cell lines. HeLa (ATCC CCL-2) epithelial human carcinoma cells were cultured in Dulbecco’s Modified Eagle’s Media GlutaMAX (DMEM, Gibco) supplemented with 10% v/v heat inactivated fetal calf serum (FCS, Gibco). THP-1 monocyte precursor cells were maintained in Roswell Park Memorial Institute (RPMI) with 200 mM Glutamax supplemented with 10% v/v FCS. All cells were maintained in 5% CO_2_ at 37°C.

*In vitro* infections were performed based on the procedure described by Klein *et. al.* 2017 (54). For HeLa infections, 5x10^4^ cells/well were seeded in 24 well plates (Corning Costar) and incubated at 37°C, 5% CO_2_ for 18-24 hours. The cells were infected with T3SS1 induced bacterial subculture at a MOI of 100 and incubated for 10 min under cell growth conditions. The cells were then washed three times with Hank’s Buffered Salt Solution (HBSS) followed by a 20 min incubation in fresh media. The media was replaced with DMEM containing 10% v/v FCS and 100 mg/ml gentamicin (Pfizer), followed by a 30min incubation. The end of this step marked the 1-hour post infection (h.p.i) timepoint. For the 5 h.p.i and 24 h.p.i, after 1 h.p.i, the culture media was replaced with DMEM supplemented with 10% v/v FCS and 10 mg/ml gentamicin. At each timepoint, the host cells were lysed with sterile 0.2% w/v sodium deoxycholate solution, and the lysates were collected for CFU estimation.

For THP-1 cells, 5x10^5^ cells/well were differentiated with 10^-8^ M of phorbol 12-myristate 13-acetate (PMA) for 72 hours at 37°C, 5% CO_2_ in 24 well plates. The remaining experimental conditions were kept the same as HeLa infection except for bacterial MOI which was 20 for these assays.

Bacterial quantification data was visualized using ggplot2 v v3.4.4 package (41) in RStudio v22.12.0 (42). To determine statistical significance levels, one way ANOVA with Dunnett’s multiple comparison tests were performed using GraphPad Prism v10.0 Mac OS X, GraphPad Software, Boston, Massachusetts USA, www.graphpad.com. For determining statistical significance levels between AUSMDU00068404 variants in THP-1 macrophages, unpaired t-test with Welch’s correction was performed using the same software.

## Results

### Multiple lineages of *S*. Panama were found to be circulating in Australia over two decades

To explore the global population of *S*. Panama and to place the Australian isolates in a broader global context, a maximum likelihood (ML) phylogeny was initially inferred with a total of 165 isolates, comprising 90 Australian and 75 selected publicly available genomes (Fig. S1). Two isolates from the USA (SRR3047883 and SRR2968115, both ST3091) and one Australian isolate (AUSMDU00038310 [ST48]) clustered on a long branch and were not considered further. The global phylogeny was thus inferred from 162 *S.* Panama isolates including 89 Australian isolates from the current study and 73 publicly available *S*. Panama genomes (Fig. 1A, Table S1, Fig. S2). All *S.* Panama isolates either belonged to ST48 or were single locus variants of ST48 (Table S1, Fig. S3). Population clustering was assessed using BAPS (39, 40). Four BAPS lineages were identified in the global *S*. Panama phylogeny (Fig. 1A). The resulting population structure was consistent with the recent findings from Pulford et al. with the Australian isolates clustering together in the phylogeny (22), forming two sub-clades represented by BAPS 3 and 4 (Fig 1). BAPS 1 consisted of isolates from several countries, while BAPS 2 was associated with isolates from the United Kingdom and Ireland. One isolate from Thailand clustered in the BAPS 4 isolates that was largely comprised of isolates from this study. A total of 89.89% (80/89) of the Australian isolates belonged to either BAPS 3 or BAPS 4, with 20% (16/80) of these forming part of BAPS 4, and the remaining 80% (64/80) clustered in BAPS 3.

**Figure 1.**
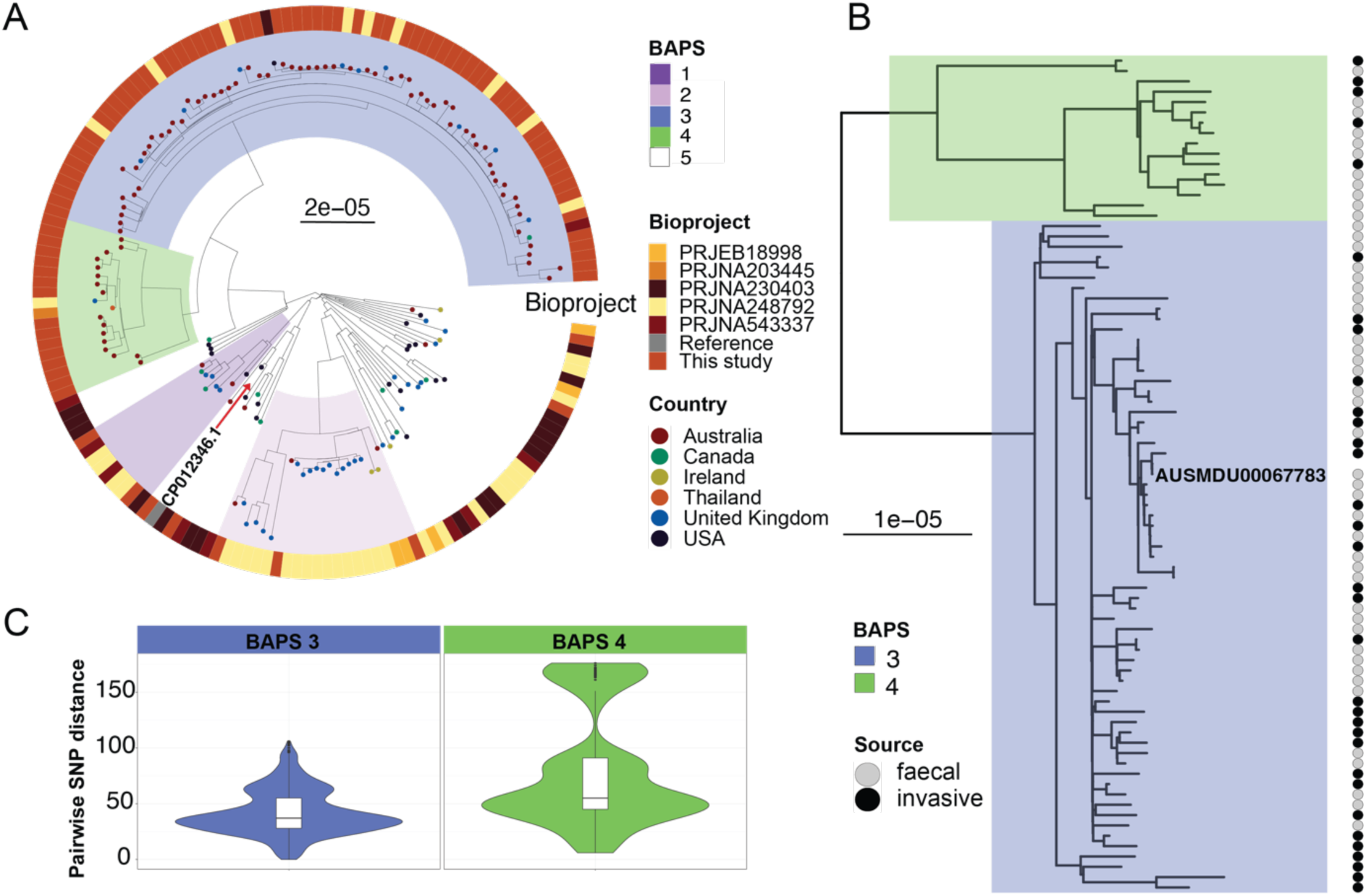
Maximum likelihood phylogenies of *Salmonella* Panama isolates from Australia and pairwise SNP distances. (**A**) Maximum likelihood (ML) tree of 162 *S.* Panama isolates including both Australian and publicly available contextual isolates. The reference was CP012346 (highlighted with red arrow). BAPS lineages were determined using RHierBAPS and are shown as coloured blocks on the tree. Tree tips indicate if the isolates were curated from public BioProjects or are from this study. The circular heatmap shows the country of isolation of the samples, as indicated by the inset legend. Scale indicates SNPs substitutions per unit branch length. (B) Local ML phylogeny of 80 Australian isolates belonging to BAPS groups 3 and 4 were mapped to the local reference AUSMDU00067783. The linked heatmap shows the distribution of faecal and invasive isolates and the coloured blocks of the tree highlight the two BAPS groups (C) Pairwise SNP distances of the BAPS lineages 3 and 4 were visualised as violin plots with box plot inset in each plot showing the median and interquartile range of the pairwise SNP distances.

In order to have higher resolution of the Australian *S.* Panama population, a ML phylogeny was inferred with the 80 Australian isolates belonging to BAPS 3 and BAPS 4, the two Australian sub-lineages of the C4 lineage previously identified (22), with a local Australian isolate, AUSMDU00067783, used as the reference strain (Fig. 1B). Invasive isolates, defined as those collected from blood samples, were present in both lineages, with the respective proportions being 43.75% (28/64) of isolates in BAPS 3 and 31.25% (5/16) of isolates in BAPS 4. Invasive isolates were not observed to be limited to either lineage, determined using two-tailed Fischer’s exact test (p=0.7721). Differences in the temporal span of the two lineages circulating in Australian were detected, BAPS 4 was comprised of *S.* Panama isolates collected between 2000 and 2012. In contrast, BAPS 3 isolates spanned the entire collection window, from 2000 to 2019, with 54.69% (35/64) of isolates collected from 2013 onwards suggestive that this lineage has replaced BAPS 4 (Table S1). Pairwise SNP distances varied between the two lineages with a wider distribution found in the older BAPS 4 lineage (median 55 pairwise SNPS, interquartile range 45-91) (Fig 1C). BAPS 3 isolates were highly clonal with limited pairwise SNPs (medium 37 SNPS, interquartile range 28-55) despite the two-decade time span.

### Multidrug resistance (MDR) is highly prevalent in Australian *S*. Panama population

AMR was common in the Australian isolates with known AMR mechanisms detected to at least one drug class in 67.5% of isolates (54/80) (Fig. 2, Table S1). Further, MDR profiles were found in 56.25% (36/64) of BAPS 3 isolates and 62.5% (10/16) of BAPS 4 isolates. No AMR mechanisms were detected that confer resistance to third-generation cephalosporins (3GCs). Of note, resistance mechanisms to azithromycin (*mphA*) and colistin (*mcr 1.1*) were detected in an invasive isolate, AUSMDU00067711 (BAPS 3). AUSMDU00067711 was inferred to be resistant to seven additional drug classes and susceptible to 3GCs and carbapenems based on the identified resistome. Limited AMR determinants against ciprofloxacin were found, with the *qnrS1* gene, associated with reduced susceptibility (59), identified in only two BAPS 3 isolates, AUSMDU00067711 and AUSMDU00066509. No point mutations in the quinolone resistance determining regions (QRDRs) were identified in any of the Australian *S*. Panama isolates.

**Figure 2.**
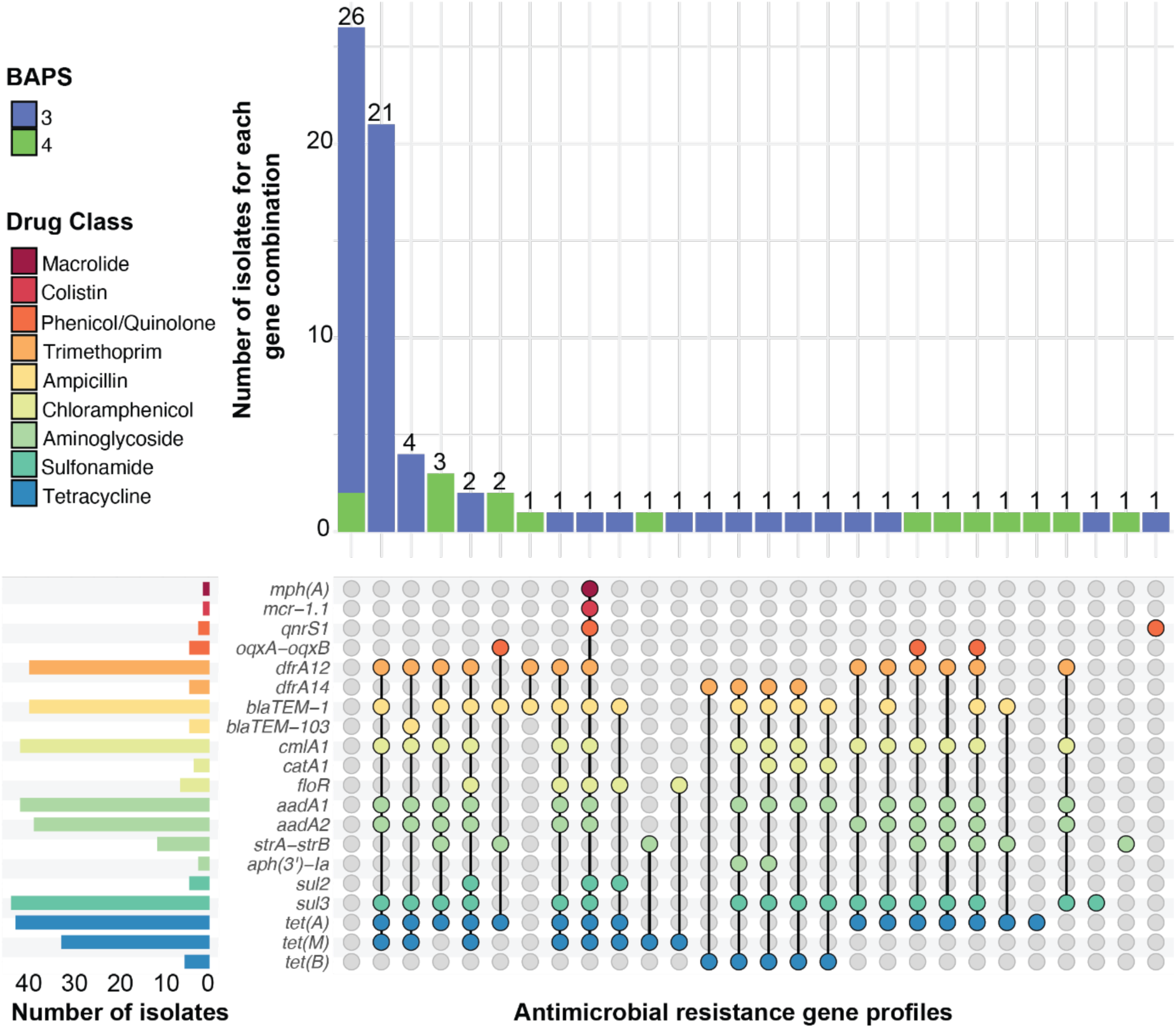
Upset plot demonstrating the antimicrobial resistance (AMR) gene profiles found in Australian *S*. Panama isolates. The y-axis bars show the total number of isolates harbouring a particular resistance gene. The stripe colours in the body of the plot correspond to the drug classes shown in the index, to which the respective genes belong. The stacked x-axis bars denote the total number of isolates from BAPS lineages 3 and 4 which have the specific combination of AMR genes, shown by the connected dots in the plot body.

Multidrug resistance in the Australian *S*. Panama isolates was common with inferred resistance to ampicillin, chloramphenicol, streptomycin, co-trimoxazole and tetracycline found in 46.25% (37/80) of the Australian BAPS 3 and BAPS 4 lineages (Fig 2). The most common AMR profile was *blaTEM-1* (ampicillin resistance), *cmlA1* (chloramphenicol resistance), *aadA1-aadA2* (streptomycin resistance), *dfrA12* and *sul3* (co-trimoxazole resistance) and *tet(A)* (tetracycline resistance), that was detected in 26.25% (21/80) of the total isolates, all of which were BAPS 3. Genes detected at lower frequencies that differed between the two populations were *tet(B)* (tetracycline) and *blaTEM-103* (ampicillin) in BAPS 3 while 12 BAPS 4 isolates carried *strA-strB* genes (streptomycin).

### Characterisation of mobile genetic elements mediating AMR in Australian *S*. Panama

The prevalence of plasmids in the *S.* Panama data was first explored by screening plasmid replicons in the draft assemblies. In BAPS 3, a total of eight different Inc types were detected, with IncFIA and IncFIIB most frequently detected in 37.5% (24/64) of BAPS 3 isolates (Fig. 3A). Further, the presence of the IncFIA and IncFIIB replicons coincided with the MDR profile of *dfrA12-blaTEM-1-cmlA1-aadA1-aadA2-sul3-tet(A)-tet(B)* in 20/24 isolates (Table S1). In BAPS 4 there were fewer plasmid replicon types detected, with only five Inc types observed across two isolates. Thus, the AMR profiles in BAPS 4 did not co-occur with plasmids replicons and suggested that the resistance genes observed in this lineage may be chromosomal (Fig 3, Table S1).

**Figure 3:**
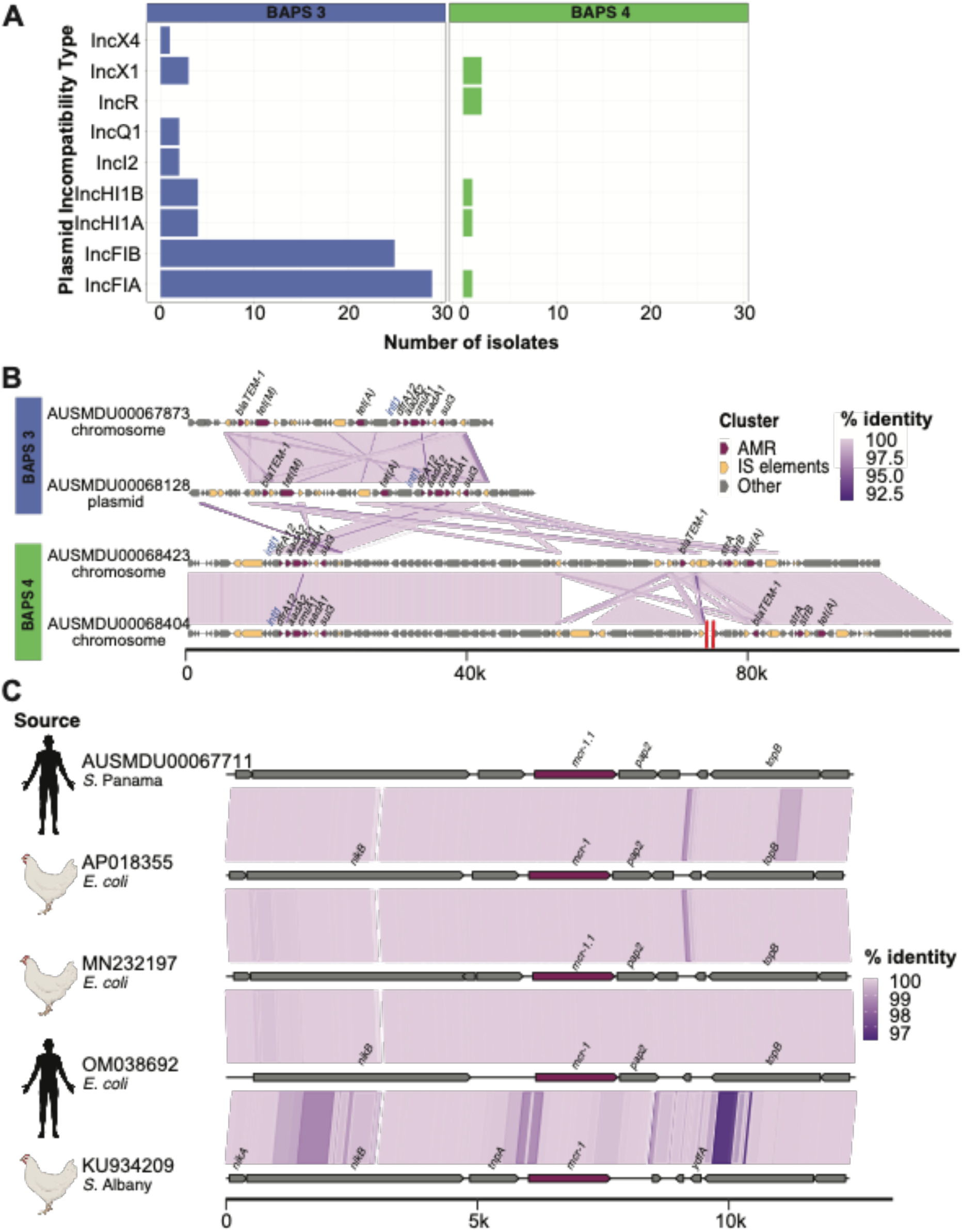
Characterisation of plasmid-mediated AMR in the Australian *S*. Panama. (A) In each panel for BAPS lineages 3 or 4, the x-axis indicates the total number of isolates carrying each plasmid Inc type shown in the y-axis. (B) The AMR genes are indicated in dark red while the insertion sequences (ISs) and transposases are shown in yellow. The purple blocks between the sequences indicate the degree of identity between the gene regions across the isolates. The red double line in the AUSMDU00068404 sequence indicates the deletion of a 405 Kb non-identical gene region to fit the sequence to a comparable scale with the other sequences. The scale at the bottom shows the genome scale in kilobases. (C) Comparison between the regions encoding *mcr-1* (shown in dark red) and its flanking genes in four IncI2 plasmids. The purple links between the sequences indicate the percentage identity between the gene regions in the isolates. The scale at the bottom shows the genome scale in kilobases. Source panel on the left shows the associated host for each isolate, with the icons retrieved from https://app.biorender.com/biorender-templates.

We investigated the complete genomes of selected MDR isolates from BAPS 3 and BAPS 4 to characterise the carriage of AMR mechanisms in the *S.* Panama isolates (Fig. 3B). Plasmid mediated MDR was limited to a single BAPS 3 isolate, AUSMDU00068128, that had two plasmid replicons detected IncFIA and IncFIIB. The AMR genes were determined to be integrated into the chromosome of the remaining three BAPS 3 isolates. The AMR genes in all isolates were carried across multiple cassettes, each cassette flanked by insertion sequences (IS) and transposon elements. The *dfrA12-aadA2-cmlA1-aadA1-sul3* cassette was common across all the isolates, sharing high sequence identity. The *intI1* coding region flanked the *dfrA12* gene that suggests that the genes are located on a Class 1 integron. The *blaTEM-1* and *tet(A)* genes occurred separately.

### Plasmid encoding *mcr-1.1* in AUSMDU00067711 suggests possible zoonotic link

Macrolide and colistin resistance genes were detected in AUSMDU00067711 (Fig 2, Table S1). Except for *mcr-1.1* (colistin resistance), all other AMR mechanisms including *mphA* (macrolide resistance) were found to be integrated into the chromosome. The *mcr-1.1* gene was carried on a 67 Kb IncI2 plasmid (Fig. S4). To determine if this was a novel plasmid or if it had been detected elsewhere, we explored publicly available data identifying highly homologous plasmids from three *E. coli* isolates collected from a human sample and two poultry samples (Accessions: OM038692.1, MN232197.1, AP018355.1) and one from *S*. Albany (KU934209.1) collected from poultry (Fig 3B). All the parent isolates for these reference plasmids were from South-East Asia. The *mcr-1.1* gene region was conserved between the AUSMDU00067711 plasmid and the reference plasmids (Fig. 3C) suggesting that this plasmid is moving between species and serovars of *Salmonella* and different hosts.

### *S*. Panama enters and replicates within macrophages but shows limited entry in epithelial cells

To investigate if the BAPS 3 and BAPS 4 lineages vary in terms of intracellular replication *in vitro*, we infected HeLa epithelial cells and differentiated THP-1 macrophage-like cells with the selected *S*. Panama isolates (Fig. 4) We further wanted to identify if the ability to infect and replicate within host cells vary between closely matched faecal and invasive isolate pairs. Only two out of seven *S*. Panama isolates entered epithelial cells, AUSMDU00068128 (BAPS 3, invasive and MDR) and AUSMDU00068423 (BAPS 4, faecal and MDR) (Fig. 4A, 4B and 4C). While AUSMDU00068128 (0.63 ± 0.28 %) showed comparable levels of epithelial invasion to *S.* Typhimurium SL1344 (1.36 ± 0.58 %), its corresponding matched faecal isolate AUSMDU00067873 did not enter HeLa cells. The entry of AUSMDU00068423 (0.069 ± 0.006 %) was significantly lower (p=0.001) than *S*. Typhimurium SL1344 (Fig. 4A), while its corresponding invasive isolate AUSMDU00068404 did not exhibit entry into epithelial cells. Over 5 hours post infection (h.p.i) and 24 h.p.i, AUSMDU00068128 replicated to significantly higher colony forming units (CFU) (2.5 x 10^5^ ± 7.1 x10^4^ CFU/ml at 5 h.p.i; 1.6 x 10^6^ ± 1.6 x 10^5^ CFU/ml at 24 h.p.i) than *S*. Typhimurium SL1344 (p=0.02 at 5 h.p.i; p<0.0001 at 24 h.p.i), while the CFU of AUSMDU00068423 (2.8 x 10^4^ ± 5.1 x 10^3^ CFU/ml at 5 h.p.i; 5.3 x 10^5^ ± 6.6 x 10^4^ CFU/ml at 24 h.p.i) was not significantly different to *S*. Typhimurium SL1344 (1.2 x 10^5^ ± 1.8 x 10^4^ CFU/ml at 5 h.p.i; 5.9 x 10^5^ ± 1.2 x 10^5^ CFU/ml at 24 h.p.i). Despite having significantly lower percentage of internalisation at 1 h.p.i in comparison with *S*. Typhimurium SL1344 (Fig. 4C) (p=0.01), AUSMDU00068423 had the highest replication rate over of 24 h.p.i in HeLa epithelial cells (73.10 ± 16.01-fold, p=0.0003) (Fig. 4B). The remaining matched isolate pair from BAPS 3, AUSMDU00067783 (faecal and no AMR genes) and AUSMDU00067801 (invasive and no AMR genes) did not enter HeLa cells.

**Figure 4:**
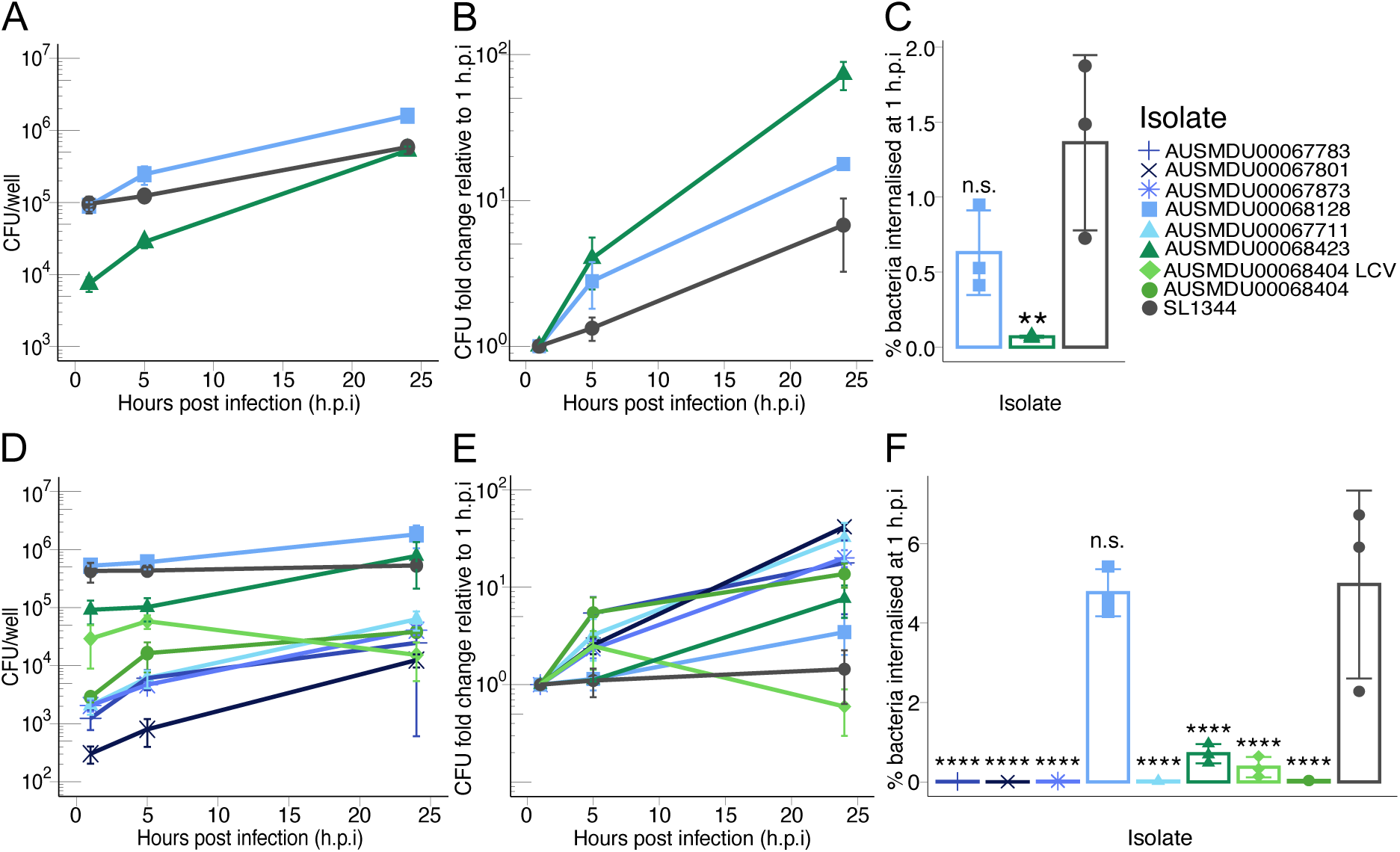
*S*. Panama shows strain-specific features of interaction with both epithelial and macrophage-like human cell lines. HeLa cells (A, B, C) and differentiated THP-1 cells (D, E, F) were infected with *S*. Panama or *S*. Typhimurium at MOI 100 and 20 respectively. (A, D) Each point indicates mean CFU/well per isolate at the three timepoints. (B, E) Each point indicates mean fold change in bacterial CFU/well of each sample with respect to 1 h.p.i. (C, F) Bars indicate mean percentage of CFU/well internalised at 1 h.p.i with respect to inoculum CFU for each isolate. The symbols indicate individual data points of the respective parameter for each sample. All experiments were performed in technical duplicates for three independent biological replicates per isolate. Error bars indicate +/- standard deviation of biological replicates. Statistical significance was determined by one-way ANOVA with Dunnett’s comparisons test at 95% confidence interval. ** indicates p<0.01 and **** indicates p<0.0001. MOI multiplicity of infection, h.p.i hours post infection, CFU colony-forming units.

All the *S*. Panama isolates were taken up by THP-1 macrophage-like cells, however levels of uptake were isolate specific. Except for AUSMDU00068128 (BAPS 3, invasive and MDR), uptake of the other *S*. Panama isolates was significantly lower than *S.* Typhimurium SL1344 (p<0.0001) (Fig. 4D and 4F). In concordance with the patterns seen in HeLa cells, AUSMDU00068128 showed the highest internalisation among the *S*. Panama isolates (4.77 ± 0.59 %), followed by AUSMDU00068423 (0.37 ± 0.26 %). At 24 h.p.i, all isolates except AUSMDU00068128 (BAPS 3), AUSMDU00068423 (BAPS 4) and AUSMDU00068404 (BAPS 4) showed significant intracellular replication, while intracellular SL1344 levels were constant (Fig. 4E, Table S2). AUSMDU00067801 (BAPS 3) showed the lowest internalisation (0.0033 ± 0.0012%), but the highest fold change of replication over 24 h.p.i (41.91 ± 3.89-fold).

Interestingly, AUSMDU00068404 (BAPS 4, invasive and MDR) showed variations with regards to colony morphology, with a large colony variant (LCV) co-occurring with medium sized colonies. Neither variant showed entry into epithelial cells. In THP-1 cells, the AUSMDU00068404 LCV showed slightly higher but non-significant uptake (0.033 ± 0.0019%) than its other colony morphology variant, AUSMDU00068404 (0.019 ± 0.0052%) (p=0.15). Both AUSMDU00068404 variants showed comparable CFU/ml at 24 h.p.i.

### Genomic comparison of closely related faecal and invasive isolate pairs that differed in phenotype

From the host cell internalisation experiments, two pairs of closely related faecal and invasive isolates showed phenotypic difference between isolates in each pair. The first matched pair was BAPS 3 MDR isolates AUSMDU00067783 (faecal) and AUSMDU00068128 (invasive). The second matched pair was BAPS 4 MDR isolates AUSMDU00068423 (faecal) and AUSMDU00068404 (invasive). To identify genomic content that differs between the matched isolates, and potentially identify the genetic basis for the observed differences in phenotype, we used both reference-based and pangenome approaches.

The matched BAPS 3 isolates were found to be highly genetically similar. AUSMDU00067783 only had two SNPs compared to AUSMDU00068128, one in a non-coding region and a missense mutation in the RNA chaperone *hfq,* and one deletion in a non-coding region. This pair of isolates shared 4459 genes and differed by four genes, three in AUSMDU00067873, while the remaining gene encodes for a hypothetical protein present in AUSMDU00068128 (Table S3). Additional efforts to characterise the hypothetical protein unique to AUSMDU00068128 using NCBI blastp identified it as the phosphate starvation-inducible protein PhoH. Further, interrogation of the NCBI Conserved Domains Database identified the PRK12567 super family (putative monovalent cation/H+ antiporter subunit B).

The matched BAPS 4 isolates had more genetic diversity detected compared to the BAPS 3 isolates. A total of 58 nucleotide differences were detected between the two isolates, including three deletions, 14 synonymous mutations and 28 missense mutations detected in coding regions (Table S4). The majority of the SNP mutations were in metabolic genes and of note was the C260A mutation (Ala87Asp) in the *Salmonella* pathogenicity island (SPI) 1 type III secretion system invasion protein gene *iagB*. Pangenome analysis found 4367 core genes and 47 accessory genes that differed between the isolates (11 in AUSMDU00068404 and 36 in AUSMDU00068423. These accessory genes consist of regulatory genes, metabolic genes, IS elements and hypothetical coding regions (Table S4).

## Discussion

*S*. Panama is an emerging iNTS serovar that has been associated with foodborne outbreaks. This study provides evidence for the establishment and continued circulation of different geographical lineages of *S.* Panama (21–23, 28). We identified two sub-lineages of *S*. Panama detected in Australia, belonging to the previously described Asia/Oceania lineage. The Australian isolates were predominantly acquired by cases while travelling to these regions. Hence, international travel is a key epidemiological risk factor for *S.* Panama infections in Australia, in particular to South-East Asia. As such, these two lineages responsible for *S.* Panama in Victoria over two-decades (2000-2020) likely reflects the *S.* Panama circulating in the broader surrounding geographical region (22). BAPS 3 has been the dominant sub-lineage in the Australian population, appearing to have displaced BAPS 4 from 2013 onwards. The unbiassed approach of sampling all *S.* Panama received in Victoria suggests that BAPS 3 isolates may also be the predominant in South-East Asia and Oceania during this period.

Similar to isolates reported from Asia, both Australian sub-lineages were associated with MDR to older antimicrobials, with >50% of the isolates possessing genetic determinants for resistance (22, 23, 27, 28). MDR in *S.* Panama is common in the Oceania and South-East Asian regions, consistent with studies of other NTS serovars from the region and reflecting likely introduction from returned travellers (30, 60–63). Of note, the MDR profile observed in this study was not restricted to plasmids, with our analyses of the completed genomes finding integration of these AMR genes into the chromosome of some isolates, movement supported by independently mobile cassettes. This suggested that a selective pressure, favouring maintenance of MDR through chromosomal integration, is acting on the *S.* Panama population. Sulfonamide resistance was mediated by *sul3* which is not frequently detected in NTS (64–66) and was mobilised in to the chromosome on a class 1 integron, which also carried *dfrA12*, *cmlA1*, and *aadA1* and *aadA2* (Genbank Accession EF051037.1). This mobile element has been previously identified in other NTS serovars, including *S.* Rissen that were isolated from pork and unknown food products in Portugal (67). The presence of cross-species plasmids and diversity of plasmid replicon genes observed in the study population demonstrates that *S.* Panama is readily able to acquire and retain AMR-containing mobile elements.

The movement of plasmids between members of the Enterobacterales has been well-established (68, 69). Here an Australian isolate of *S.* Panama was shown to possess a single IncI plasmid harbouring a colistin resistance gene, *mcr1.1*, that was homologous to IncI plasmids detected in other serovars of *Salmonella* and *E. coli* collected from poultry in the South-East Asian and Oceanic regions. While directionality of plasmid transfer is unable to be inferred from the current data, this does provide additional evidence for the ongoing emergence of drug resistance underpinned by plasmids moving within and between bacterial species with broad host ranges, including livestock animals. A key challenge to identifying the spread of AMR via the food supply chain is the lack of *S.* Panama data from food sources and livestock. To further explore dissemination of AMR through food products, livestock and humans, future studies comprising a ‘One Health’ approach to sampling strategies are required (70).

This study provides the first data on the host-pathogen interactions of *S*. Panama. We examined the ability of multiple faecal and invasive isolate pairs from both sub-lineages to enter and replicate in both epithelial and macrophage-like cell lines and observed higher replication of *S*. Panama inside macrophages. These *in vitro* infection experiments demonstrated that individual isolates interact differently with host cells. Importantly, no phenotype was clearly associated with the invasive isolates. Rather, all isolates demonstrated a capacity for replication in macrophage-like cells and this may be a trait that reflects the potential for development of iNTS. Studies have shown that invasive variants of *S*. Typhimurium, such as *S*. Typhimurium ST313, or monophasic variants of the same serovar, show increased survival in macrophages possibly linked to reduced programmed cell death activation in the host cell (15, 61). *S.* Typhimurium ST313 also induces reduced secretion of proinflammatory cytokines which may aid in its immune evasion and dissemination inside the human host (15). Additionally, another iNTS, *S*. Dublin that is commonly found in cattle, has been shown to induce less cell death of bovine macrophages (13). Interestingly, the *S.* Panama isolates that entered THP-1 cells with greater efficiency did not replicate as much as those with lower internalisation. This may a link between bacterial load and host cell response. Characterisation of host cell death induced by *S*. Panama and its regulation of host immune response will provide valuable insights into mechanisms of iNTS infections.

A 2018 outbreak study from Taiwan by Feng et al. showed high internalisation and survival of *S*. Panama isolates in macrophages *in vitro* compared to *S*. Typhimurium. However, they only investigated two isolates (21). In this study, only AUSMDU00068128, an invasive isolate from BAPS 3 entered both cell types in comparable numbers to *S*. Typhimurium and survived better over time. But corresponding faecal isolate AUSMDU00067873 did not enter epithelial cells and showed restricted entry into macrophages. Genetic comparison between AUSMDU00067873 and AUSMDU00068128 revealed minor SNP and gene content differences, which shows they are closely related despite their different phenotypes in HeLa cells. The four ORFs divergently encoded by these isolates do not provide insight into the genetic basis for these different phenotypes. However, it is possible that the missense mutation within Hfq could influence the post-transcriptional regulation of SPI-1 (71–73). Future studies exploring the proteome of these two isolates may provide further insight into their distinct host-pathogen interactions. AUSMDU00068423, a faecal isolate from BAPS 4 entered HeLa cells at much lower numbers but showed similar replication to *S*. Typhimurium, while its corresponding invasive isolate did not invade HeLa cells. This BAPS 4 isolate pair had a higher number of SNPs and gene content compared to the divergent BAPS 3 pair. These differences are likely because these isolates do not share the same immediate common ancestor in the phylogeny. While a missense point mutation was detected in *iagB*, its encoded effector IagB acts as an accessory SPI-1 gene and is not essential for SPI-1 regulation (74). Given these observations, it would be pivotal to investigate the regulation of *S*. Panama effectors that are responsible for host cell entry and intracellular bacterial replication to shed more light on its pathogenesis.

This study provides detailed insights into the evolution and genomic epidemiology of clinical isolates of *S.* Panama circulating in Victoria, Australia over a two-decade period. This unbiassed sampling approach is a key strength of the study, enabling insights into *S.* Panama in the region over an extended time. While source of isolation (blood or faecal) was available for the *S.* Panama data, additional health information including co-morbidities, of the patients was not available which is a limitation of the study. Given the source of collection did not correlate with phenotypic data, host factors are likely to play an important role in the ability of *S.* Panama to cause iNTS, with all faecal isolates potentially able to cause invasive infections. This is consistent with other iNTS serovars (6). This research provides a foundation to understand the pathogenesis of a highly invasive but understudied NTS serovar that is prevalent in Australia and worldwide.

## Acknowledgements

DJI is supported by an NHMRC Investigator Grant (GNT1195210). Research in HJNs laboratory was funded by the NHMRC (GNT2010841). ST is supported by University of Melbourne Graduate Researcher Scholarship.

## References

1. Gal-Mor O, Boyle EC, Grassl GA. Same species, different diseases: how and why typhoidal and non-typhoidal *Salmonella enterica* serovars differ. Front Microbiol. 2014;5:391.

2. Majowicz SE, Musto J, Scallan E, Angulo FJ, Kirk M, O’Brien SJ, et al. The global burden of nontyphoidal *Salmonella* gastroenteritis. Clin Infect Dis. 2010;50(6):882–9.

3. Tennant SM, Diallo S, Levy H, Livio S, Sow SO, Tapia M, et al. Identification by PCR of non-typhoidal *Salmonella enterica* serovars associated with invasive infections among febrile patients in Mali. PLoS Negl Trop Dis. 2010;4(3):e621.

4. Crump JA, Sjolund-Karlsson M, Gordon MA, Parry CM. Epidemiology, Clinical Presentation, Laboratory Diagnosis, Antimicrobial Resistance, and Antimicrobial Management of Invasive *Salmonella* Infections. Clin Microbiol Rev. 2015;28(4):901–37.

5. Collaborators GBDN-TSID. The global burden of non-typhoidal *Salmonella* invasive disease: a systematic analysis for the Global Burden of Disease Study 2017. Lancet Infect Dis. 2019;19(12):1312–24.

6. Jones TF, Ingram LA, Cieslak PR, Vugia DJ, Tobin-D’Angelo M, Hurd S, et al. Salmonellosis outcomes differ substantially by serotype. J Infect Dis. 2008;198(1):109–14.

7. Vugia DJ, Samuel M, Farley MM, Marcus R, Shiferaw B, Shallow S, et al. Invasive *Salmonella* infections in the United States, FoodNet, 1996-1999: incidence, serotype distribution, and outcome. Clin Infect Dis. 2004;38 Suppl 3:S149–56.

8. Weinberger M, Solnik-Isaac H, Shachar D, Reisfeld A, Valinsky L, Andorn N, et al. *Salmonella enterica* serotype Virchow: epidemiology, resistance patterns and molecular characterisation of an invasive *Salmonella* serotype in Israel. Clin Microbiol Infect. 2006;12(10):999–1005.

9. Langridge GC, Wain J, Nair S. Invasive Salmonellosis in Humans. EcoSal Plus. 2012;5(1).

10. Campioni F, Gomes CN, Bergamini AMM, Rodrigues DP, Tiba-Casas MR, Falcao JP. Comparison of cell invasion, macrophage survival and inflammatory cytokines profiles between *Salmonella enterica* serovars Enteritidis and Dublin from Brazil. J Appl Microbiol. 2021;130(6):2123–31.

11. Campioni F, Vilela FP, Cao G, Kastanis G, Dos Prazeres Rodrigues D, Costa RG, et al. Whole genome sequencing analyses revealed that *Salmonella enterica* serovar Dublin strains from Brazil belonged to two predominant clades. Sci Rep. 2022;12(1):10555.

12. Fong WY, Canals R, Predeus AV, Perez-Sepulveda B, Wenner N, Lacharme-Lora L, et al. Genome-wide fitness analysis identifies genes required for *in vitro* growth and macrophage infection by African and global epidemic pathovariants of *Salmonella enterica* Enteritidis. Microb Genom. 2023;9(5).

13. Huang K, Fresno AH, Skov S, Olsen JE. Dynamics and Outcome of Macrophage Interaction Between *Salmonella* Gallinarum, *Salmonella* Typhimurium, and *Salmonella* Dublin and Macrophages From Chicken and Cattle. Front Cell Infect Microbiol. 2019;9:420.

14. Mohammed M, Le Hello S, Leekitcharoenphon P, Hendriksen R. The invasome of *Salmonella* Dublin as revealed by whole genome sequencing. BMC Infect Dis. 2017;17(1):544.

15. Ramachandran G, Perkins DJ, Schmidlein PJ, Tulapurkar ME, Tennant SM. Invasive *Salmonella* Typhimurium ST313 with naturally attenuated flagellin elicits reduced inflammation and replicates within macrophages. PLoS Negl Trop Dis. 2015;9(1):e3394.

16. Van Puyvelde S, Pickard D, Vandelannoote K, Heinz E, Barbe B, de Block T, et al. An African *Salmonella* Typhimurium ST313 sublineage with extensive drug-resistance and signatures of host adaptation. Nat Commun. 2019;10(1):4280.

17. Kingsley RA, Msefula CL, Thomson NR, Kariuki S, Holt KE, Gordon MA, et al. Epidemic multiple drug resistant *Salmonella* Typhimurium causing invasive disease in sub-Saharan Africa have a distinct genotype. Genome Res. 2009;19(12):2279–87.

18. Kumwenda B, Canals R, Predeus AV, Zhu X, Kroger C, Pulford C, et al. *Salmonella enterica* serovar Typhimurium ST313 sublineage 2.2 has emerged in Malawi with a characteristic gene expression signature and a fitness advantage. Microlife. 2024;5:uqae005.

19. Martins IM, Seribelli AA, Machado Ribeiro TR, da Silva P, Lustri BC, Hernandes RT, et al. Invasive non-typhoidal *Salmonella* (iNTS) aminoglycoside-resistant ST313 isolates feature unique pathogenic mechanisms to reach the bloodstream. Infect Genet Evol. 2023;116:105519.

20. Pulford CV, Perez-Sepulveda BM, Canals R, Bevington JA, Bengtsson RJ, Wenner N, et al. Stepwise evolution of *Salmonella* Typhimurium ST313 causing bloodstream infection in Africa. Nat Microbiol. 2021;6(3):327–38.

21. Feng Y, Chen CL, Chang YJ, Li YH, Chiou CS, Su LH, et al. Microbiological and genomic investigations of invasive *Salmonella enterica* serovar Panama from a large outbreak in Taiwan. J Formos Med Assoc. 2022;121(3):660–9.

22. Pulford CV, Perez-Sepulveda BM, Ingle DJ, Bengtsson RJ, Bennett RJ, Rodwell EV, et al. Global diversity and evolution of *Salmonella* Panama, an understudied serovar causing gastrointestinal and invasive disease worldwide: a genomic epidemiology study. bioRxiv. 2024:2024.02.09.579599.

23. Pulford CV, Perez-Sepulveda BM, Rodwell EV, Weill FX, Baker KS, Hinton JCD. *Salmonella enterica* Serovar Panama, an Understudied Serovar Responsible for Extraintestinal Salmonellosis Worldwide. Infect Immun. 2019;87(9).

24. Authority EFS, Prevention ECfD, Control. The European Union summary report on trends and sources of zoonoses, zoonotic agents and food-borne outbreaks in 2012. EFSA Journal. 2014;12(2):3547.

25. Bonardi S, Bruini I, Bolzoni L, Cozzolino P, Pierantoni M, Brindani F, et al. Assessment of *Salmonella* survival in dry-cured Italian salami. Int J Food Microbiol. 2017;262:99–106.

26. Lee JA. Recent trends in human salmonellosis in England and Wales: the epidemiology of prevalent serotypes other than *Salmonella typhimurium*. J Hyg (Lond). 1974;72(2):185–95.

27. Lee HY, Yang YJ, Su LH, Hsu CH, Fu YM, Chiu CH. Genotyping and antimicrobial susceptibility of *Salmonella enterica* serotype Panama isolated in Taiwan. J Microbiol Immunol Infect. 2008;41(6):507–12.

28. Matsushita S, Kawamura M, Takahashi M, Yokoyama K, Konishi N, Yanagawa Y, et al. Serovar-distribution and drug-resistance of *Salmonella* strains isolated from domestic and imported cases during 1995-1999 in Tokyo. Kansenshogaku Zasshi. 2001;75(2):116–23.

29. Sherry NL, Horan KA, Ballard SA, Gonҫalves da Silva A, Gorrie CL, Schultz MB, et al. An ISO-certified genomics workflow for identification and surveillance of antimicrobial resistance. Nat Commun. 2023;14(1):60.

30. Williamson DA, Lane CR, Easton M, Valcanis M, Strachan J, Veitch MG, et al. Increasing Antimicrobial Resistance in Nontyphoidal *Salmonella* Isolates in Australia from 1979 to 2015. Antimicrob Agents Chemother. 2018;62(2).

31. Sia CM, Baines SL, Valcanis M, Lee DYJ, Goncalves da Silva A, Ballard SA, et al. Genomic diversity of antimicrobial resistance in non-typhoidal *Salmonella* in Victoria, Australia. Microb Genom. 2021;7(12).

32. Wood DE, Salzberg SL. Kraken: ultrafast metagenomic sequence classification using exact alignments. Genome Biol. 2014;15(3):R46.

33. Bankevich A, Nurk S, Antipov D, Gurevich AA, Dvorkin M, Kulikov AS, et al. SPAdes: a new genome assembly algorithm and its applications to single-cell sequencing. J Comput Biol. 2012;19(5):455–77.

34. Yoshida CE, Kruczkiewicz P, Laing CR, Lingohr EJ, Gannon VP, Nash JH, Taboada EN. The Salmonella *In Silico* Typing Resource (SISTR): An Open Web-Accessible Tool for Rapidly Typing and Subtyping Draft *Salmonella* Genome Assemblies. PLoS One. 2016;11(1):e0147101.

35. Arndt D, Grant JR, Marcu A, Sajed T, Pon A, Liang Y, Wishart DS. PHASTER: a better, faster version of the PHAST phage search tool. Nucleic Acids Res. 2016;44(W1):W16–21.

36. Zhou Y, Liang Y, Lynch KH, Dennis JJ, Wishart DS. PHAST: a fast phage search tool. Nucleic Acids Res. 2011;39(Web Server issue):W347–52.

37. Croucher NJ, Page AJ, Connor TR, Delaney AJ, Keane JA, Bentley SD, et al. Rapid phylogenetic analysis of large samples of recombinant bacterial whole genome sequences using Gubbins. Nucleic Acids Res. 2015;43(3):e15.

38. Minh BQ, Schmidt HA, Chernomor O, Schrempf D, Woodhams MD, von Haeseler A, Lanfear R. IQ-TREE 2: New Models and Efficient Methods for Phylogenetic Inference in the Genomic Era. Mol Biol Evol. 2020;37(5):1530–4.

39. Cheng L, Connor TR, Siren J, Aanensen DM, Corander J. Hierarchical and spatially explicit clustering of DNA sequences with BAPS software. Mol Biol Evol. 2013;30(5):1224–8.

40. Tonkin-Hill G, Lees JA, Bentley SD, Frost SDW, Corander J. RhierBAPS: An R implementation of the population clustering algorithm hierBAPS. Wellcome Open Res. 2018;3:93.

41. Wickham H. ggplot2. Wiley interdisciplinary reviews: computational statistics. 2011;3(2):180–5.

42. Team R. RStudio: Integrated Development for R. RStudio, PBC, Boston, MA URL. 2020.

43. Feldgarden M, Brover V, Gonzalez-Escalona N, Frye JG, Haendiges J, Haft DH, et al. AMRFinderPlus and the Reference Gene Catalog facilitate examination of the genomic links among antimicrobial resistance, stress response, and virulence. 2021;11(1):1–9.

44. Lex A, Gehlenborg N, Strobelt H, Vuillemot R, Pfister H. UpSet: Visualization of Intersecting Sets. IEEE Trans Vis Comput Graph. 2014;20(12):1983–92.

45. Krassowski M. ComplexUpset. ComplexUpset. 2020.

46. Carattoli A, Hasman H. PlasmidFinder and *In Silico* pMLST: Identification and Typing of Plasmid Replicons in Whole-Genome Sequencing (WGS). Methods Mol Biol. 2020;2075:285–94.

47. Carattoli A, Zankari E, Garcia-Fernandez A, Voldby Larsen M, Lund O, Villa L, et al. *In silico* detection and typing of plasmids using PlasmidFinder and plasmid multilocus sequence typing. Antimicrob Agents Chemother. 2014;58(7):3895–903.

48. Baines SL, da Silva AG, Carter GP, Jennison A, Rathnayake I, Graham RM, et al. Complete microbial genomes for public health in Australia and the Southwest Pacific. Microb Genom. 2020;6(12).

49. Wick RR, Holt KE. Polypolish: Short-read polishing of long-read bacterial genome assemblies. PLoS Comput Biol. 2022;18(1):e1009802.

50. Schwengers O, Jelonek L, Dieckmann MA, Beyvers S, Blom J, Goesmann A. Bakta: rapid and standardized annotation of bacterial genomes via alignment-free sequence identification. Microb Genom. 2021;7(11).

51. Sullivan MJ, Petty NK, Beatson SA. Easyfig: a genome comparison visualizer. Bioinformatics. 2011;27(7):1009–10.

52. Hackl T, Ankenbrand M, van Adrichem B. gggenomes: A Grammar of Graphics for Comparative Genomics. R package version 0.9.12.9000 2023 [Available from: https://github.com/thackl/gggenomes.

53. Alikhan NF, Petty NK, Ben Zakour NL, Beatson SA. BLAST Ring Image Generator (BRIG): simple prokaryote genome comparisons. BMC Genomics. 2011;12:402.

54. Klein JA, Powers TR, Knodler LA. Measurement of *Salmonella enterica* Internalization and Vacuole Lysis in Epithelial Cells. Methods Mol Biol. 2017;1519:285–96.

55. Seemann T. Prokka: rapid prokaryotic genome annotation. Bioinformatics. 2014;30(14):2068–9.

56. Tonkin-Hill G, MacAlasdair N, Ruis C, Weimann A, Horesh G, Lees JA, et al. Producing polished prokaryotic pangenomes with the Panaroo pipeline. Genome biology. 2020;21(1):1–21.

57. Wang J, Chitsaz F, Derbyshire MK, Gonzales NR, Gwadz M, Lu S, et al. The conserved domain database in 2023. Nucleic Acids Res. 2023;51(D1):D384–D8.

58. Wheeler DL, Barrett T, Benson DA, Bryant SH, Canese K, Chetvernin V, et al. Database resources of the National Center for Biotechnology Information. Nucleic Acids Res. 2007;35(Database issue):D5–12.

59. Vien le TM, Abuoun M, Morrison V, Thomson N, Campbell JI, Woodward MJ, et al. Differential phenotypic and genotypic characteristics of *qnrS1*-harboring plasmids carried by hospital and community commensal enterobacteria. Antimicrob Agents Chemother. 2011;55(4):1798–802.

60. Chung The H, Pham P, Thanh Tuyen H, Vo Kim Phuong L, Phuong Yen N, Le S-NH, et al. Multidrug resistance plasmids underlie clonal expansions and international spread of *Salmonella enterica* serotype 4,[5],12,i:-ST34 in Southeast Asia. Commun Biol. 2023; 6. 10.1038/s42003-023-05365-1

61. Ingle DJ, Ambrose RL, Baines SL, Duchene S, Goncalves da Silva A, Lee DYJ, et al. Evolutionary dynamics of multidrug resistant *Salmonella enterica* serovar 4,[5],12:i:-in Australia. Nat Commun. 2021;12(1):4786.

62. Sriyapai P, Pulsrikarn C, Chansiri K, Nyamniyom A, Sriyapai T. Molecular Characterization of Cephalosporin and Fluoroquinolone Resistant *Salmonella* Choleraesuis Isolated from Patients with Systemic Salmonellosis in Thailand. Antibiotics (Basel). 2021;10(7).

63. Woh PY, Yeung MPS, Goggins WB, 3rd, Lo N, Wong KT, Chow V, et al. Genomic Epidemiology of Multidrug-Resistant Nontyphoidal *Salmonella* in Young Children Hospitalized for Gastroenteritis. Microbiol Spectr. 2021;9(1):e0024821.

64. Antunes P, Machado J, Sousa JC, Peixe L. Dissemination of sulfonamide resistance genes (*sul1*, *sul2*, and *sul3*) in Portuguese *Salmonella enterica* strains and relation with integrons. Antimicrob Agents Chemother. 2005;49(2):836–9.

65. Guerra B, Junker E, Helmuth R. Incidence of the recently described sulfonamide resistance gene *sul3* among German *Salmonella enterica* strains isolated from livestock and food. Antimicrob Agents Chemother. 2004;48(7):2712–5.

66. Maka L, Mackiw E, Sciezynska H, Modzelewska M, Popowska M. Resistance to Sulfonamides and Dissemination of *sul* Genes Among *Salmonella* spp. Isolated from Food in Poland. Foodborne Pathog Dis. 2015;12(5):383–9.

67. Antunes P, Machado J, Peixe L. Dissemination of *sul3*-containing elements linked to class 1 integrons with an unusual 3’ conserved sequence region among *Salmonella* isolates. Antimicrob Agents Chemother. 2007;51(4):1545–8.

68. Kim YJ, Seo KH, Kim S, Bae S. Phylogenetic Comparison and Characterization of an *mcr-1*-Harboring Complete Plasmid Genome Isolated from Enterobacteriaceae. Microb Drug Resist. 2022;28(4):492–7.

69. Matlock W, Lipworth S, Chau KK, AbuOun M, Barker L, Kavanagh J, et al. Enterobacterales plasmid sharing amongst human bloodstream infections, livestock, wastewater, and waterway niches in Oxfordshire, UK. Elife. 2023;12.

70. Thorpe HA, Booton R, Kallonen T, Gibbon MJ, Couto N, Passet V, et al. A large-scale genomic snapshot of *Klebsiella* spp. isolates in Northern Italy reveals limited transmission between clinical and non-clinical settings. Nat Microbiol. 2022;7(12):2054–67.

71. Abdulla SZ, Kim K, Azam MS, Golubeva YA, Cakar F, Slauch JM, Vanderpool CK. Small RNAs activate *Salmonella* pathogenicity island 1 by modulating mRNA stability through the *hilD* mRNA 3′ UTR. bioRxiv. 2022:2022.09.07.507058.

72. El Mouali Y, Gaviria-Cantin T, Sanchez-Romero MA, Gibert M, Westermann AJ, Vogel J, Balsalobre C. CRP-cAMP mediates silencing of *Salmonella* virulence at the post-transcriptional level. PLoS Genet. 2018;14(6):e1007401.

73. Lopez-Garrido J, Puerta-Fernandez E, Casadesus J. A eukaryotic-like 3’ untranslated region in *Salmonella enterica hilD* mRNA. Nucleic Acids Res. 2014;42(9):5894–906.

74. Sukhan A, Kubori T, Wilson J, Galan JE. Genetic analysis of assembly of the *Salmonella enterica* serovar Typhimurium type III secretion-associated needle complex. J Bacteriol. 2001;183(4):1159–67.

