## Supplementary Data for "Genomic epidemiology and phenotypic characterisation of *Salmonella enterica* serovar Panama in Victoria, Australia"

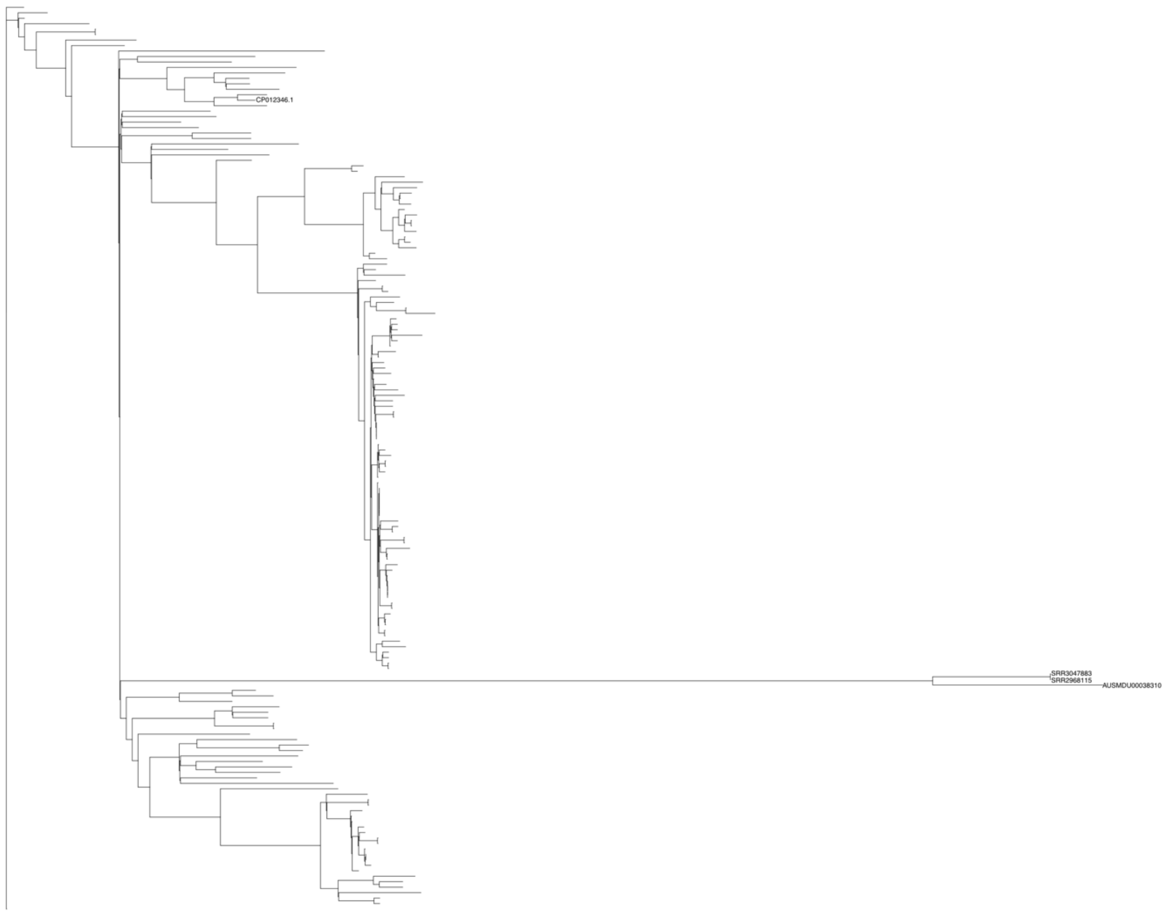


**Supplementary Figure 1: Unrooted global maximum likelihood phylogeny of all isolates**

The phylogeny was inferred from a total of 165 isolates including 90 Australian and 75 publicly available isolates, with CP012346 as the reference strain. Three isolates on the long branch, AUSMDU00038310, SRR3047883 and SRR2968115 were excluded from subsequent analyses.

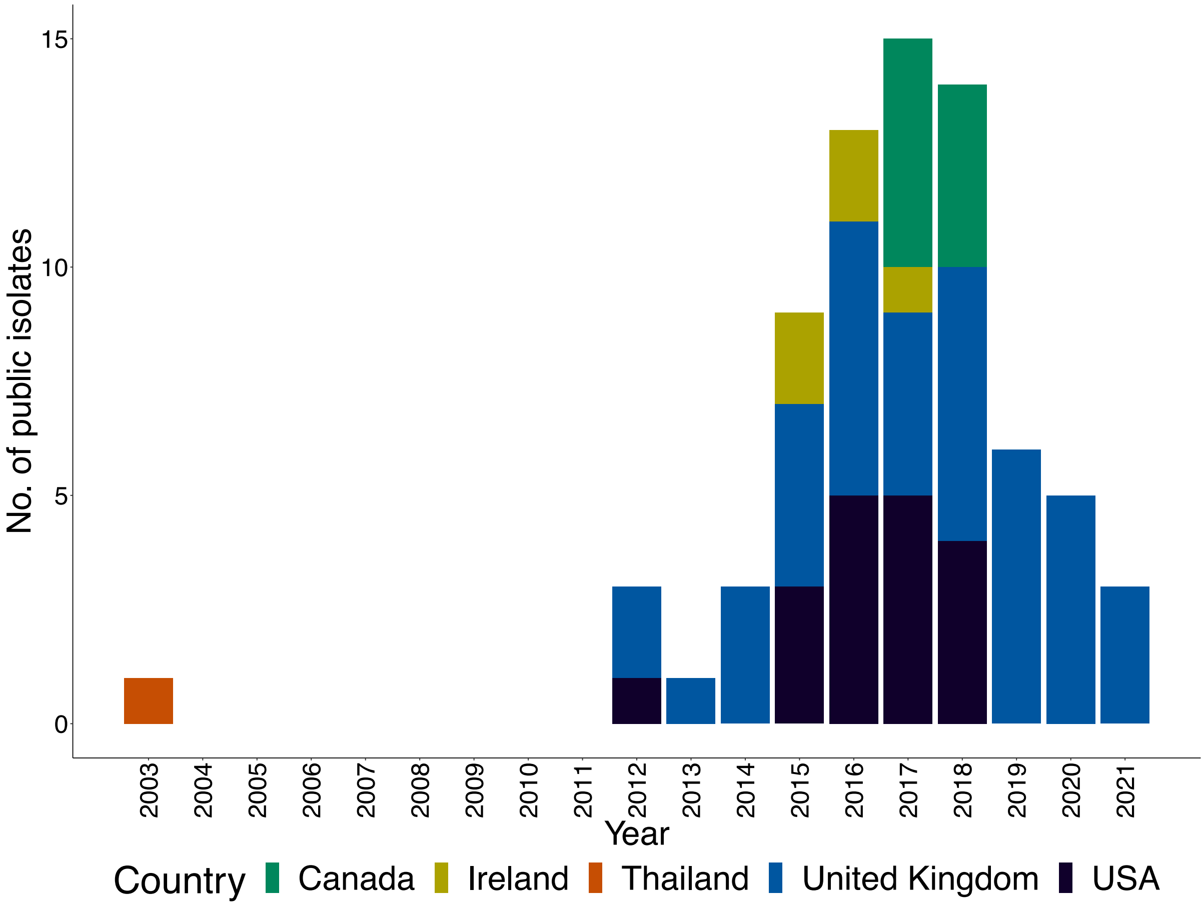


**Supplementary Figure 2: Reported year of collection and geographical site of the publicly available isolates included in the study.**
The stacked bars indicate total number of isolates that were sampled from each year from the different countries designated by the colours.

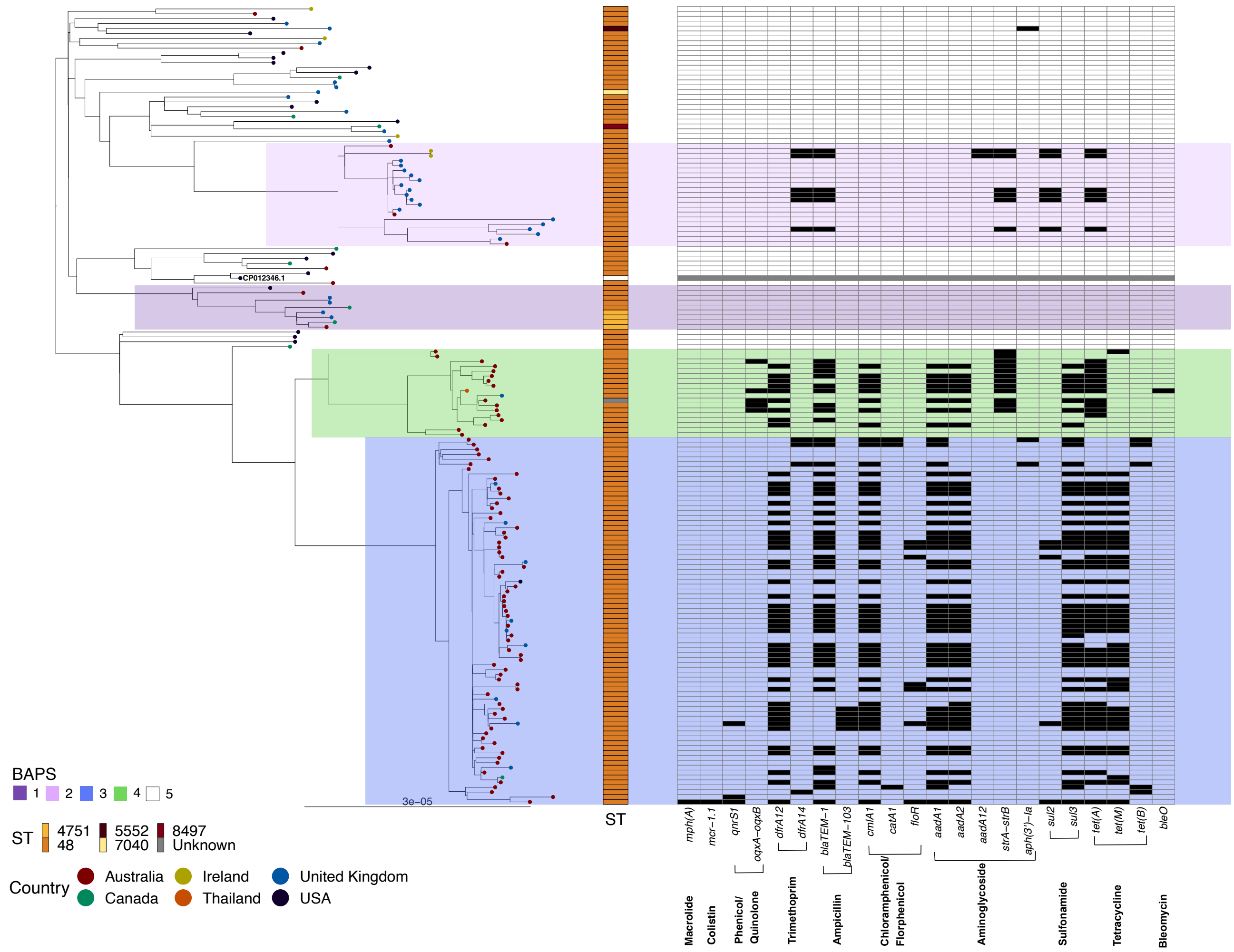


**Supplementary Figure 3: Global ML phylogeny showing the ST and AMR gene distribution**

The phylogeny was inferred from 162 isolates including 89 Australian and 73 publicly available isolates, with CP012346 as the reference strain. The highlighted clades indicate BAPS lineages. Tree tips are coloured by country of isolation of the samples. The sequence type (ST) are shown to the right of the tree. The AMR heatmap shows the genes detected in AbritAMR for the complete dataset. Scale indicates SNPs substitutions per unit branch length.

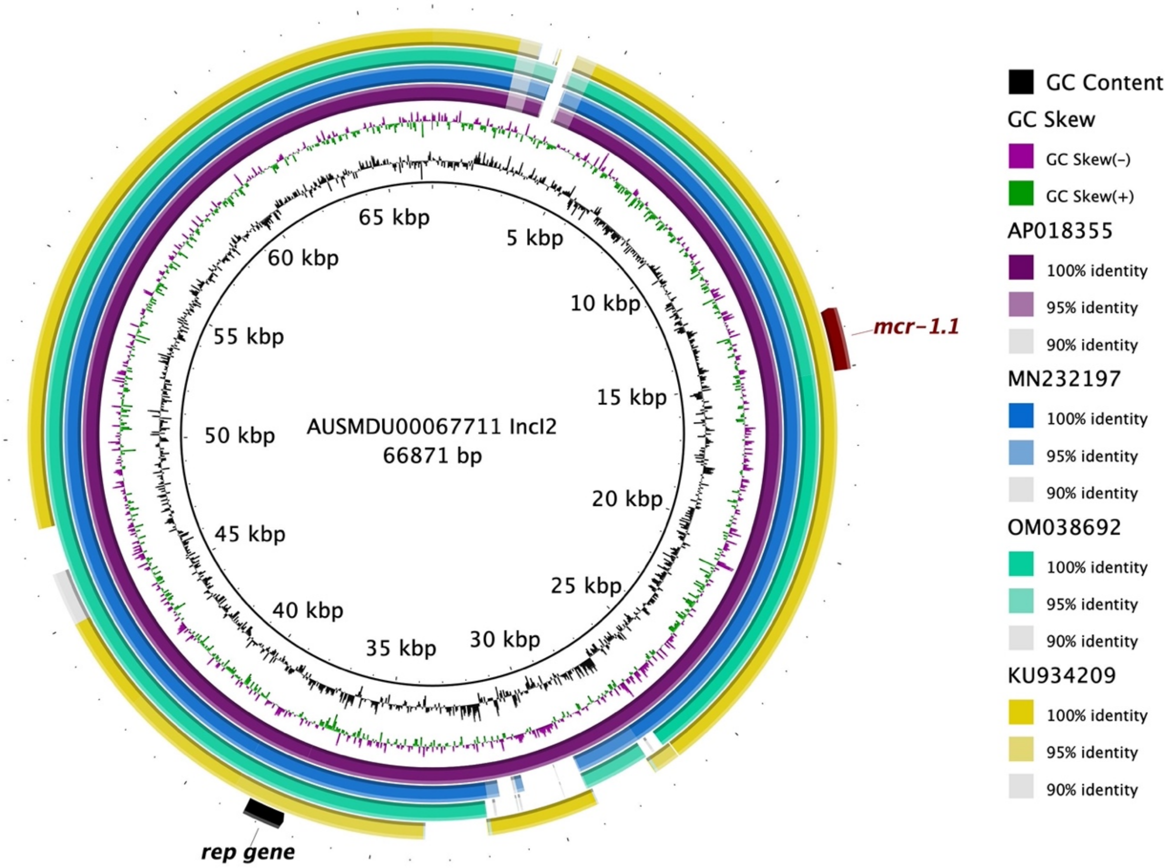


**Supplementary Figure 4: Comparison of publicly available plasmids with IncI plasmid in AUSMDU00067711**

BRIG plot demonstrating the alignment of the plasmid from the BAPS lineage 3 isolate AUSMDU00067711 harbouring the *mcr1.1* gene to public reference IncI2 plasmids from *E. coli* (Accessions: OM038692.1, MN232197.1, AP018355.1) and *S.* Albany (KU934209.1).
